# DNA Content and DNA Damage in Raw and Heat-processed Foods

**DOI:** 10.1101/2025.09.21.677616

**Authors:** Jinwoo Shin, Pawel Jaruga, Miral Dizdaroglu, Eric T. Kool

## Abstract

DNA in foods is a source of nucleotides that are salvaged by tissues as building blocks for chromosomal and mitochondrial DNA. A recent study provided preliminary evidence that high-temperature cooking damages the DNA in foods and suggested that certain forms of DNA damage can be taken up as nucleotides via metabolic salvage in cells and animals, directly incorporating genotoxic and mutagenic species into the host DNA. To assess potential risks, we surveyed DNA in 21 food ingredients, including plant- and meat-based foods in raw and roasted form. We find large variation in extractable DNA content, implying widely variable levels of consumption. Cooking resulted in greatly elevated levels of oxidative and deaminated DNA damage in nearly all foods as indicated by 8-oxo-dG and dU nucleotides; with up to 250-fold increases. Studies of human cell lines found that incubation with these damaged nucleosides resulted in cytotoxicity and increased DNA double-strand break levels.

## INTRODUCTION

Nucleotide salvage pathways exist in mammals to secure DNA and RNA components from cellular turnover and from the diet and convert them into useful building blocks for new cellular nucleic acids as well as energy sources.^1-5^ Although human cells have active *de novo* nucleotide synthesis biochemical pathways,^1^ salvage pathways exist in parallel as a means of obtaining DNA building blocks at a lower energetic cost. Among biomolecules that comprise the human diet, a large amount of research has gone into carbohydrates, proteins, and lipids, which are used as energy sources or as components for intracellular biomolecular synthesis.^6-10^ However, nucleic acid salvage and reuse from the diet has been less well studied, particularly for the DNA component of foods.^5, 9-14^

The steps of nucleotide salvage for DNA initially involve digestion of the biopolymer by nucleases in the stomach and intestine into short oligomers and ultimately mononucleotides (Fig. 1).^11, 12^ 5’-Nucleotidase activity in the cell membrane of these tissues then removes the 5’-phosphate from nucleotides, promoting cellular uptake of the uncharged mononucleosides.^15^ Once inside the cell, pyrimidine deoxynucleosides are subject to the simplest salvage pathway, as they can, in some cases, be directly phosphorylated by kinases to produce deoxynucleoside triphosphates (dNTPs) as substrates for DNA polymerases.^2, 3^ Purine deoxynucleoside salvage is more complex, and involves depurination of the base from deoxyribose, followed by re-assembly during ribonucleotide synthesis and reductive conversion to the 2’-deoxynucleoside.^3^ Subsequent 5’-phosphorylation steps, carried out by nucleoside/tide kinases, convert the nucleosides into 5’-triphosphate derivatives to act as building blocks for chromosomal and mitochondrial DNA synthesis.

**Figure 1.**
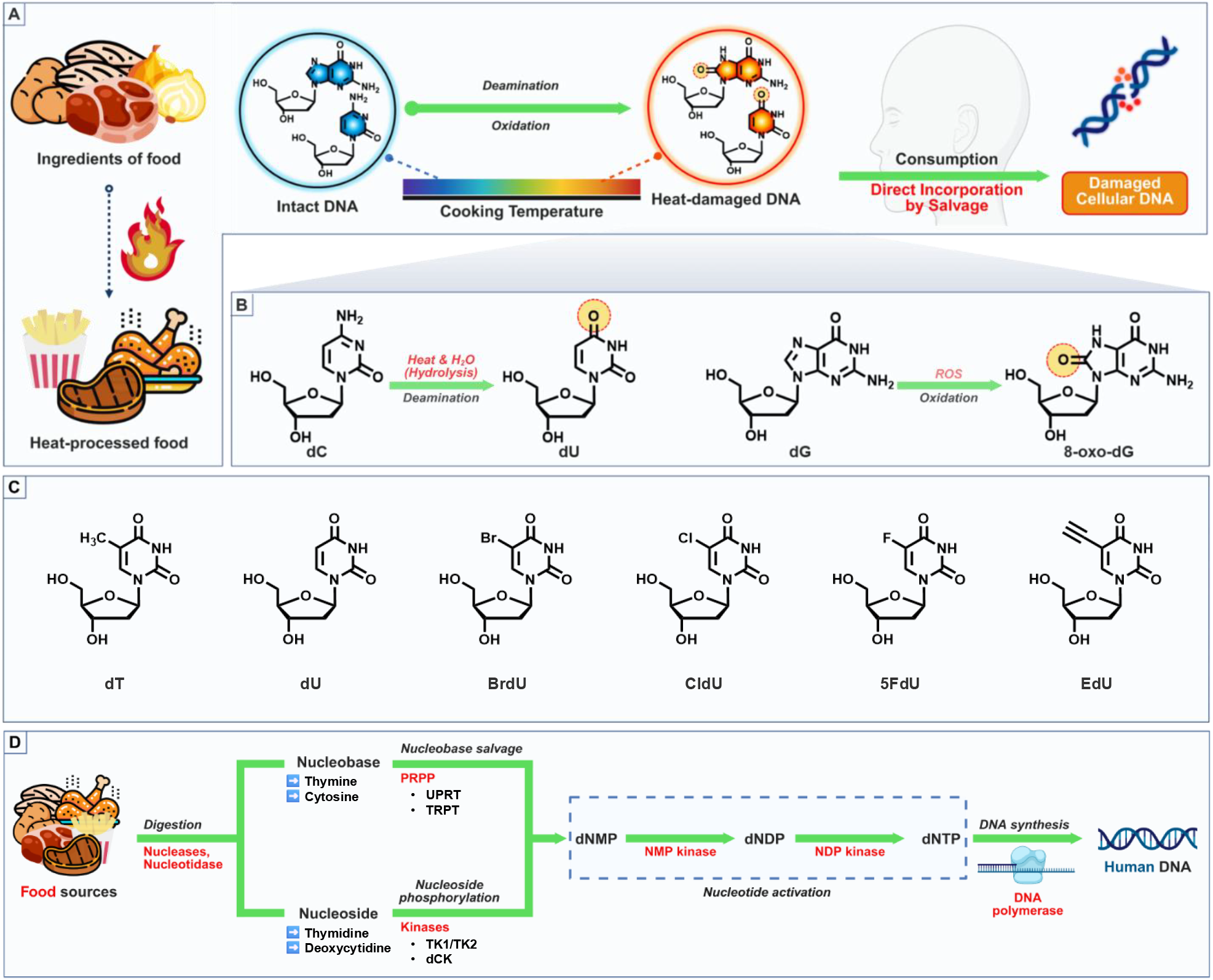
(A) Schematic illustration of proposed genotoxic and mutagenic pathway involving the direct incorporation of heat-damaged dietary DNA components into cellular DNA. (B) Known mechanisms for heat-induced DNA damage, highlighting representative deaminative and oxidative lesions. (C) Chemical structures of nucleosides differentialy modified at the 5-position, all of which are previously documented to be incorporated into the DNA via the salvage pathway. Shown are a canonical nucleoside (dT), a damaged nucleoside (dU), and modified nucleoside analogs (BrdU, CldU, 5FdU, and EdU). (D) Schematic illustration of the pathways of pyrimidine salvage from the diet. Abbreviations: UPRT, uracil phosphoribosyltransferase; TRPT, thymine phosphoribosyltransferase; TK1/TK2, thymidine kinases 1/2; dCK, deoxycytidine kinase; dNMP, deoxynucleoside monophosphate; NDP, deoxynucleoside diphosphate; dNTP, nucleoside triphosphate.

Although nucleotide salvage pathways are well studied, surprisingly little is known about the salvage of DNA from the diet.^10, 11, 13, 14^ Human diets contain relatively large amounts of DNA that can be subject to salvage (Figs. 1A and 1B),^14, 16-18^ with estimates of up to 1 g / day depending on diet.^16^ Although ribonucleotides and RNA are in greater abundance and are also subject to salvage,^5^ their pathway to DNA synthesis is considerably more complex than for the pyrimidine deoxynucleotide components of DNA, which is relatively direct as outlined above. It is not yet clear to what extent *de novo* synthesis versus salvage accounts for the composition of DNA in humans, but certain rapidly dividing tissues, such as intestinal epithelial cells and hematopoietic stem cells, appear to be largely dependent on salvage for DNA synthesis.^4, 19^ In addition, isotopic labeling experiments have measured up to half of the DNA pyrimidine composition in tumor cell lines resulting from salvage.^2, 20^ Mononucleotides have been documented as nutrients, particularly in infant formula and in mammalian diets.^4, 9, 21^ Another important documentation of deoxynucleotide uptake in humans is the mechanism of action of certain nucleotide drugs (*vide infra*). Aside from measurements of the digestion of DNA to deoxynucleotides,^11, 12^ little data exists on tracking the salvage of DNA from the diet in humans or animal models. Questions that remain include: to what degree does deoxynucleotide salvage occur in humans relative to *de novo* synthesis? How does this ratio vary with age and with tissue type? And finally, does salvage occur with noncanonical deoxynucleotides from the diet, and if so, to what extent?

While the salvage pathways evolved to capture and recycle canonical nucleotides (dA, dT, dG, and dC as well as ribonucleotides) for DNA synthesis, the enzymes in the pathway are known to be leaky, and can also take up modified nucleotides to a significant extent. An important example of this is the salvage of synthetically modified therapeutic nucleosides, a strategy that is well established for anticancer drugs in the clinic.^22, 23^ For example, 5-Fluoro-pyrimidines as free bases and as deoxynucleosides (*e*.*g*., 5-fluorodeoxyuridine (5FdU)) are important anticancer agents, and have been documented to be incorporated into the DNA of tumor cells.^24^ A second example is the antitumor drug gemcitabine, another pyrimidine deoxynucleotide analog that is incorporated into cellular DNA after salvage.^2, 25^

In addition to therapeutic applications, nucleotide salvage is also widely employed as an important research tool for labeling and tracking DNA synthesis in human cell studies. Metabolic uptake of modified pyrimidine deoxynucleosides is commonly used as a marker for new DNA synthesis; nucleosides used for this purpose include 5-bromo-dU, 5-chloro-dU, and 5-ethynyl-dU.^26-28^ These are structurally similar to thymidine (Fig. 1C) which enables them to pass through the canonical deoxypyrimidine salvage, phosphorylation and DNA synthesis pathways, and are typically incubated with cells at 10 mmol/L to 250 mmol/L for 1 h to 4 h prior to cell fixation and fluorescence imaging of the labeled newly synthesized DNA. This application, combined with the knowledge of therapeutic nucleoside drugs and their incorporation into DNA, makes it clear that the mammalian cellular salvage pathway is imperfectly selective and can accept structures different from those of the four canonical deoxynucleosides.

Given the leakiness of the nucleotide salvage pathway, we ask whether noncanonical DNA nucleotides that occur in the diet might also find their way into human DNA. It has long been known that heating in air increases the levels of damaged nucleotides in DNA *in vitro*, including deaminative damage (chiefly 2’-deoxyuridine, dU) and oxidative damage (such as 8-oxo-2’-deoxyguanosine, 8-oxo-dG) (Fig. 1B).^29, 30^ These two damaged nucleotides, when they arise in human cellular DNA, are sources of genotoxicity and mutagenesis that can lead to pathologies including cancer.^31, 32^ Combining these observations, it is plausible that cooking foods in air might damage the food DNA and that after digestion, damaged nucleotides might participate to some extent in nucleotide salvage metabolic pathways and enter cellular DNA (Fig. 1D). To begin to address this issue, a preliminary study in our laboratory measured levels of damaged deoxynucleotides in the DNA of pork, beef, and potato in raw and cooked form, and reported increases of up to 20-fold in DNA damage for the meats after roasting.^12^ Interestingly, the levels of damage were considerably lower in potato relative to the meats, suggesting the possible existence of DNA-protective factors in that plant-based food. The study also documented incubating cultured human cells with several damaged nucleosides, and presented evidence that some of them induce DNA damage and DNA repair responses in the cells, particularly for selected forms of damaged pyrimidines that closely resemble canonical ones. Feeding mice diets supplemented with dU or with DNA containing dU resulted in increases of up to 15-fold in dU content in the genomic DNA of the small intestine.^12^ Overall, those early experiments suggested the possibility that damaged nucleotides in foods that arise from high-temperature cooking have the potential to participate in the salvage pathway and thus present a possible risk in human diets, at least for high-level exposure.

In this new study, we ask how general the effect of DNA damage in heat-processed foods is. First, it is unclear how much DNA exists in different food sources; this is important because even if damage occurs, it may exist in only small quantities if there is little DNA content to begin with. To address this, here we measure DNA content in a broad range of food sources, and we then examine whether deaminative and oxidative DNA damage resulting from high temperature exposure is different in, for example, plant relative to animal tissues, to identify foods with especially high DNA content and elevated damage content. Third, we evaluate how varied preparation methods might variably affect the levels of damage in foods with the highest damage exposures; and finally, we compare the genotoxicity of two of the most common lesions in DNA, dU and 8-oxo-dG, in human cell lines. The results provide new insights into nutritional components and potential risks in human diets.

## MATERIALS AND METHODS

### Materials

Food ingredients were purchased from Safeway (Menlo Park, CA, USA). Easy-DNA gDNA purification kit was purchased from Thermo Fisher Scientific (K180001). Damaged nucleosides used for cellular experiments were purchased from Sigma-Aldrich. Phospho-Histone H2AX (Ser 139) Rabbit mAb was purchased from Cell Signaling Technology (9718S). This was visualized with Goat anti-Rabbit IgG secondary antibody, Alexa Fluor Plus 647 (Invitrogen, A32733). 2′-Deoxyuridine-^13^C_9_,^15^N_2_ and 8-oxo-dG-^15^N_5_ were purchased from Cambridge Isotope Laboratories, Inc. (Tewksbury, MA, USA).

### DNA extraction from food samples

Food ingredients and detailed sources are listed in the Supporting Data file. Prior to cooking, all food ingredients were cut into cubes (1 cm × 1 cm). Foods were prepared using one of five methods: (1) raw (uncooked control), (2) boiling in water at approximately 100 ℃ for 25 min, (3) sous vide preparation, with vacuum-sealing and cooking at 60 ℃ for 1 h, (4) frying in a pan containing vegetable oil maintained at 200 ℃ for 20 min, turning the food every 5 min for even cooking, and (5) roasting for 15 min in a standard kitchen oven preheated to 220 ℃.

### DNA content

DNA was extracted using an Easy-DNA gDNA purification kit (Thermo Fisher Scientific, K180001). Briefly, food samples were blotted with paper towels to remove excess moisture, chopped into small pieces, and incubated with Solution A and Solution B (provided in the kit) following the manufacturer’s protocol. After extraction with chloroform and subsequent centrifugation, DNA was precipitated and purified by ethanol precipitation according to the manufacturer’s instructions. DNA yields were calculated based on the total amount of DNA (mg) obtained relative to the initial mass of food samples prepared. Yields are based on food wet weights rather than dried.

### Quantification of DNA damage in extracted DNA

Quantitative analysis of DNA damage (specifically 8-oxo-dG and dU) in extracted DNAs was performed. Aliquots of 50 μg from three independently isolated DNA samples per food type were subjected to analysis by LC-MS/MS. LC-MS/MS was utilized for quantifying 8-oxo-dG and dU with internal standards including 8-oxo-dG-^15^N_5_ and dU-^13^C_9_, ^15^N_2_. Hydrolysis conditions and detailed analytical procedures for releasing and quantifying the damage products as nucleosides have been described previously.^33^ The mass-to-charge (m/z) transitions used were m/z 229 → m/z 113 for dU, m/z 240 → m/z 119 for dU-^13^C_9_,^15^N_2_, m/z 284→m/z 168 for 8-oxo-dG, and m/z 289→m/z 173 for 8-oxo-dG-^15^N_5_. Measurements were performed in triplicate and reported as average ± standard deviation (S.D.). Additional methodological details are included in the Supporting file.

### Immunostaining of γ-H2AX

DSBs in cells were assessed via immunostaining of γ-H2AX, following the manufacturer’s instructions. Human colorectal cancer cells (HCT116) were cultured in 1× McCoy’s 5A medium supplemented with 10% fetal bovine serum (FBS) and 1% penicillin-streptomycin. Cells were maintained in a humidified incubator at 37 ℃ with 5% CO2 and passaged upon reaching approximately 80% confluency. For DNA damage induction, cells were treated with 200 µmol/L or 1 mmol/L of the damaged nucleosides 8-oxo-dG and dU. After incubation, the medium was removed and cells were washed three times with PBS. Fixation was carried out using 4% methanol-free formaldehyde in PBS for 10 minutes at room temperature, followed by three additional PBS washes. Cells were then permeabilized with 0.5% Triton X-100 in PBS for 5 min at room temperature, washed once with PBS, and blocked with 3% BSA in PBS for 1 hour at room temperature. After blocking, cells were incubated with a γH2AX (Ser139) primary antibody diluted in PBS containing 1% BSA and 1% Triton X-100 for 24 h at 4 ℃. The antibody solution was removed, and cells were washed three times with PBS. Alexa Fluor 647-conjugated goat anti-rabbit IgG secondary antibody, diluted in PBS with 1% BSA and 1% Triton X-100, was then added and incubated for 1 hour at room temperature. After three PBS washes, nuclei were stained with 1 µg/mL Hoechst 33342 in PBS with 1 % BSA and 1 % Triton X-100 for 30 minutes at room temperature. Following staining, cells were imaged using a confocal microscope with excitation at 405 nm for Hoechst 33342 and 647 nm for Alexa Fluor 647. Image acquisition and quantification of γ-H2AX foci were performed using Fiji (ImageJ) software.

### Cell viability measurement

Cell viability was assessed using the LDH-Glo Cytotoxicity Assay (Promega) according to the manufacturer’s instructions, with slight modifications. HCT116 colorectal cancer cells were seeded at a density of 20000 cells per well in a 96-well white opaque plate (100 µL of 2 × 10^5^ cells/mL suspension per well) in 1× McCoy’s 5A medium supplemented with 10% FBS and 1% penicillin-streptomycin. Cells were incubated overnight at 37 ℃ in a 5% CO2 humidified incubator to allow for attachment and stabilization. The next day, culture medium was aspirated and replaced with 100 µL of fresh medium containing damaged nucleosides (8-oxo-dG or dU) at concentrations of (50, 100, 200, and 500) µmol/L, 1 mmol/L, or 2 mmol/L. Wells containing only medium (no cells) were included as negative controls (background), and wells seeded with cells but treated with 2 µL of 10% Triton X-100 served as maximum LDH release controls. In addition, vehicle-only control wells (cells treated with fresh medium without damaged nucleosides) were included to serve as a baseline for comparison. After 24 hours of incubation, 2 µL of culture supernatant was collected from each well. For the maximum release control, 2 µL of 10% Triton X-100 was added 10 minutes before sampling to lyse the cells. All collected samples were diluted with 48 µL of LDH Storage Buffer (25-fold dilution) and kept on ice. For further dilution to 100-fold, 25 µL of the diluted sample was mixed with 75 µL of LDH Storage Buffer. Then, 50 µL of the final sample was transferred to a new 96-well plate and combined with 50 µL of freshly prepared LDH Detection Reagent. After incubation at room temperature for 60 minutes, luminescence was measured using a microplate reader.

Cytotoxicity was calculated using the following formula:

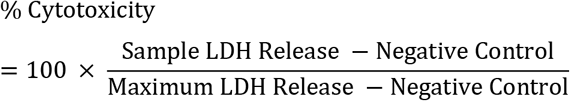

Cell viability was determined as:

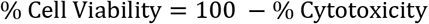

## RESULT AND DISCUSSION

### DNA content

Our long-term goal is to evaluate potential risks, if any, of damaged DNA in heat-processed foods. In principle, any risk from a specific damaged nucleotide should exist as a product of the relative level of damage per total nucleotide content multiplied by the DNA content of the food, which has the potential to vary significantly. This would represent the dietary exposure to a damaged nucleotide of interest. Thus, our initial experiments were directed to quantifying the DNA content of foods, as there is relatively little information on this in the literature. Montag and coworkers measured RNA and DNA content for a variety of raw foods after drying,^17, 18^ but we were unable to find few other general measurements, and none after cooking.

We employed a commercial kit to extract genomic DNA from 21 food ingredients (Table S1) with DNA affinity columns (see detailed materials and methods in the Supporting Data file). Foods were chosen based on a variety of plant and animal sources; we did not select fruits for this study because they are less frequently cooked than are meats and vegetables. Weighed food samples were subject to flash freezing, grinding, extraction, and isolation on DNA affinity columns. This was performed in triplicate for each food source, and DNA yields were then measured via UV absorbance (A_260_). Foods for the broad survey were measured both in raw form and after mild roasting for 15 min at 220 ℃ prior to extraction, and DNA yields were normalized as mg/gram of food. Results of DNA content measurements are given in Fig. 2 and Table 1. We term our data “extractable DNA content” rather than total DNA content to reflect the fact that these measurements of DNA content are distinct from how humans extract DNA from foods via chewing and digestion, and yields in both cases likely vary depending on the different food matrices.

**Table 1.** Extractable DNA content from various food ingredients.

| Food Ingredients | Extraction Yield ($\mu\text{g/g}$)<br>of Roasted Samples<sup>a, b</sup> | Rank<sup>c</sup> | Extraction Yield ($\mu\text{g/g}$)<br>of Raw Samples<sup>a, b</sup> | Rank<sup>c</sup> |
| --- | --- | --- | --- | --- |
| Mushroom | 4700 ± 270 | 1 | 2100 ± 20 | 1 |
| Cauliflower | 3800 ± 50 | 2 | 360 ± 8.0 | 10 |
| Garlic | 2900 ± 50 | 3 | 450 ± 20 | 7 |
| Chicken | 2400 ± 140 | 4 | 430 ± 40 | 8 |
| Beef substitute | 2300 ± 30 | 5 | 1600 ± 40 | 2 |
| Shrimp | 2100 ± 50 | 6 | 590 ± 30 | 6 |
| Salmon | 1400 ± 10 | 7 | 330 ± 10 | 12 |
| Green bean | 1300 ± 6.0 | 8 | 700 ± 10 | 5 |
| Spinach | 1300 ± 20 | 8 | 340 ± 9.0 | 11 |
| Tofu | 1200 ± 130 | 9 | 910 ± 2.0 | 3 |
| Asparagus | 1100 ± 40 | 10 | 860 ± 20 | 4 |
| Broccoli | 1000 ± 60 | 11 | 400 ± 8.0 | 9 |
| Onion | 420 ± 10 | 12 | 61 ± 2.0 | 19 |
| Carrot | 370 ± 6.0 | 13 | 81 ± 2.0 | 18 |
| White fish | 350 ± 9.0 | 14 | 260 ± 2.0 | 14 |
| Almond | 350 ± 20 | 14 | 330 ± 20 | 12 |
| Pork | 290 ± 7.0 | 15 | 230 ± 0.5 | 15 |
| Beef | 190 ± 6.0 | 16 | 230 ± 0.5 | 15 |
| Cheese | 120 ± 30 | 17 | 140 ± 0.2 | 16 |
| Bacon | 100 ± 10 | 18 | 280 ± 5.0 | 13 |
| Potato | 96 ± 0.3 | 19 | 120 ± 1.0 | 17 |

<sup>a</sup> Values shown as $\mu\text{g}$ DNA per gram of food in roasted and raw forms. <sup>b</sup> Data represent mean ± S.D. from at least three replicates. <sup>c</sup> Ranked based on extraction yields from roasted samples.

**Figure 2.**
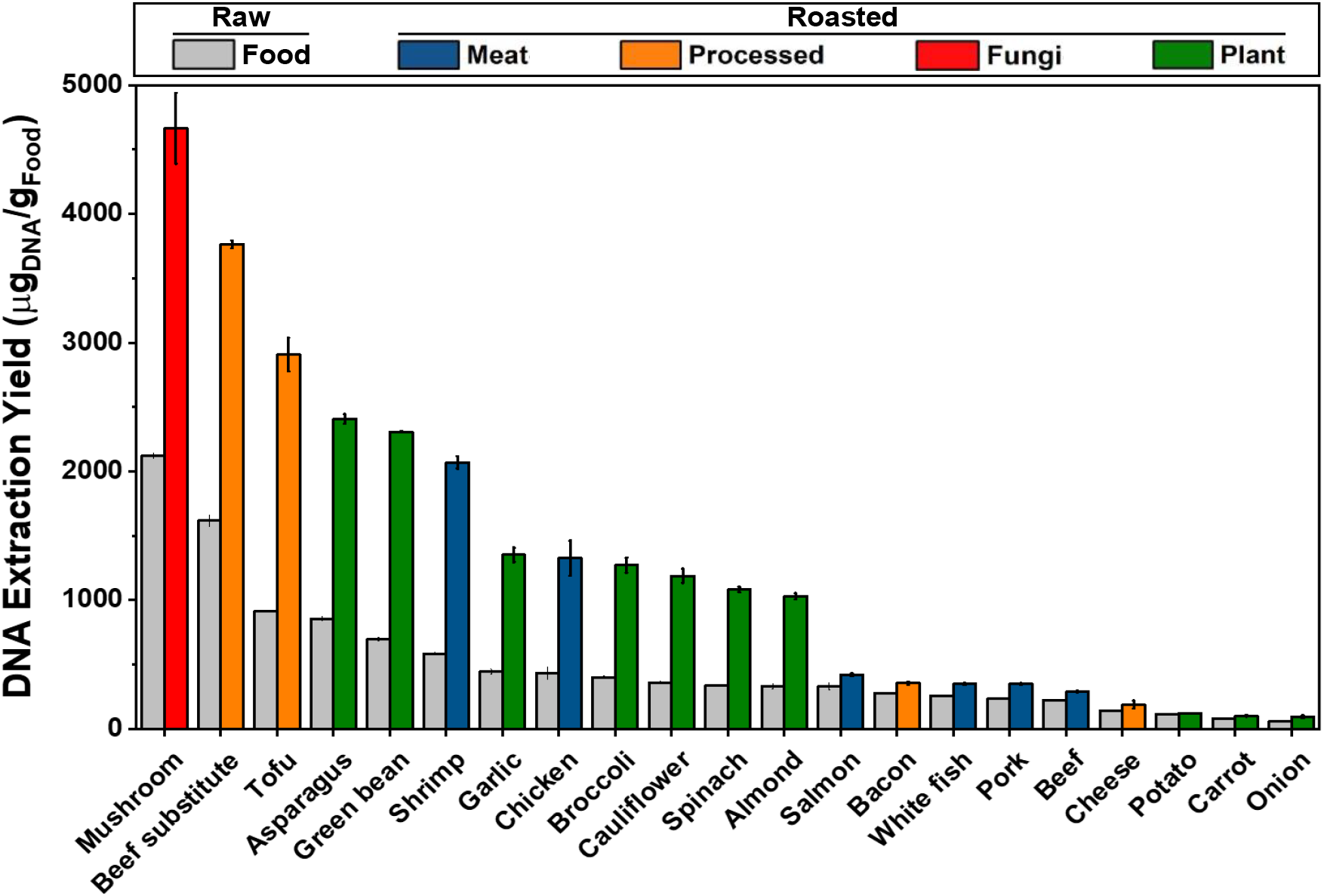
Extractable DNA content (μg of DNA per gram of food) from various food ingredients. DNA extraction yields were calculated based on the total amount of DNA obtained relative to the initial mass of each food sample. Data are presented for both raw and roasted forms, categorized as plant-based, meat-based, marine-based, or processed-derived foods. Data represent the mean ± S.D. from at least three replicates.

The data shows that extractable DNA content varies widely in different foods. We find a 35-fold variation in DNA content for the raw foods and 48-fold in roasted foods. Interestingly, roasting increases extractable DNA content by a significant factor (*ca*. 2-fold) in nearly all cases, presumably by concentrating the food (via water evaporation) and possibly also by helping to break up cell and tissue matrices more efficiently than does grinding alone. Looking at variation among foods, there appears to be no general trend with type of food; for example, we find that plant-based sources can have either very high or quite low DNA content. Interestingly, the meats tested tend toward the center of the DNA content distribution, whereas plants (and a fungus) lie at the extreme high and low ends. Notably, the three highest DNA contents are found in mushrooms, a beef-simulating meat substitute, and tofu, all commonly used in diets as replacements for meat. For raw mushrooms (the sole representative of fungi here), we found DNA content at 2 mg/g, 0.2 % of the total food content by weight. Among plants, asparagus and green beans offered the highest DNA content (700 mg/g to 860 mg/g), while potatoes and carrots yielded much lower DNA quantities, at 60mg/g to 80 mg/g. Among meats, the variation was smaller at 230 mg/g (beef) to 580 mg/g (shrimp).

### DNA damage levels and exposure

Next, we measured levels of deaminated and oxidatively damaged nucleotides in DNA extracted from these foods before (*i*.*e*., raw) and after roasting. We chose dU as a measure of deamination damage; dU is among the most common lesions in natural DNA sources,^31^ and is known to increase markedly in levels with exposure to heat.^29^ Uracil, when it forms from cytidine in genomic DNA, causes mutations, and in elevated levels leads to DNA double strand breaks (DSB) as cells attempt to repair it via base excision.^31^ Mutations resulting from deamination predominate as C>T mutations, a common signature found in tumor DNA.^34^ As a measure of oxidative damage, we evaluated 8-oxo-dG, which has been cited as the most common oxidized lesion in DNA.^32^ 8-oxo-dG in DNA is highly mutagenic lesion leading to G>T transversions and defects in its repair are known to lead to mutations that cause cancer.^32, 34^ The damaged nucleotides were quantified by LC-MS/MS following published methods^12^. After nuclease and alkaline phosphatase digestion of the extracted DNA to nucleosides measurements were performed in triplicates.

Fig. 3A shows a plot of levels of dU in the DNA of raw and cooked foods (tabulated in Table 2). For the large majority of cases (19/21), roasting markedly increased the levels of dU damage, while no increase was seen for tofu (Table 3). The high-temperature exposure increased dU frequency by up to 250-fold for some of the foods. Highest dU frequencies were seen for roasted salmon, beef, cheese, and mushrooms, at ≈400 to 950 lesions per million bases; at the highest frequencies this lesion represents ca. 0.1 % of the mass of the DNA. Lowest frequencies among the roasted foods occurred with carrot, asparagus, bacon, and potato (ca. 3-10 lesions/million bases). As expected, numbers were low for raw foods, from ≈20 dU per 10^6^ bases to undetectable levels (<2 lesions/10^6^ bases). For reference, numbers for dU lesions are reported to be similarly low for DNA in human cells, at *ca*. 1-10 per 10^6^ bases.^35^

**Table 2.** Levels of deaminative DNA damage (dU) in food ingredients.

| Food Ingredients | dU Lesions<br>/10<sup>6</sup> DNA Bases<sup>a</sup> |  | Fold-Increase<sup>b</sup> | Rank<sup>b</sup> |
| --- | --- | --- | --- | --- |
|  | Roasted<sup>c</sup> | Raw<sup>c</sup> |  |  |
| Mushroom | 160 ± 4.0 | 0.6 ± 0.2 | 250 | 1 |
| Salmon | 950 ± 42 | 15 ± 1.3 | 62 | 2 |
| Chicken | 85 ± 3.7 | 1.7 ± 0.4 | 51 | 3 |
| Onion | 98 ± 4.0 | 2.0 ± 0.4 | 49 | 4 |
| Cheese | 360 ± 34 | 8.8 ± 1.8 | 41 | 5 |
| Beef | 640 ± 130 | 17 ± 3.1 | 38 | 6 |
| Broccoli | 31 ± 1.5 | 1.4 ± 0.3 | 22 | 7 |
| White fish | 59 ± 0.4 | 3.5 ± 0.2 | 17 | 8 |
| Cauliflower | 42 ± 3.6 | 3.0 ± 0.4 | 14 | 9 |
| Spinach | 12 ± 0.6 | 0.9 ± 0.1 | 13 | 10 |
| Green bean | 13 ± 0.1 | 1.0 ± 0.04 | 13 | 11 |
| Almond | 13 ± 0.4 | 1.6 ± 0.1 | 7.8 | 12 |
| Shrimp | 110 ± 7.4 | 16 ± 0.9 | 7.2 | 13 |
| Pork | 44 ± 2.2 | 6.1 ± 1.0 | 7.2 | 14 |
| Beef substitute | 56 ± 3.6 | 13 ± 0.9 | 4.2 | 15 |
| Bacon | 2.2 ± 0.4 | 0.8 ± 0.04 | 2.7 | 16 |
| Asparagus | 8.2 ± 0.8 | 3.2 ± 0.4 | 2.6 | 17 |
| Potato | 9.0 ± 0.3 | 4.0 ± 1.0 | 2.2 | 18 |
| Tofu | 6.9 ± 0.8 | 15 ± 0.2 | 0.5 | 19 |
| Garlic | 37 ± 2.2 | *n.d.* | *n.d.* | 20 |
| Carrot | 4.6 ± 0.3 | *n.d.* | *n.d.* | 21 |

<sup>a</sup> Expressed as lesions per 10<sup>6</sup> DNA bases (roasted and raw). <sup>b</sup> Fold-increase (roasted/raw) calculated and ranked accordingly. <sup>c</sup>
Data represent mean ± S.D. from three replicates (*n* = 3).

**Table 3.** Levels of oxidative DNA damage (8-oxo-dG) in food ingredients.

| Food Ingredients | 8-oxo-dG Lesions<br>/10<sup>6</sup> DNA Bases<sup>a</sup> |  | Fold Increase<sup>b</sup> | Rank<sup>b</sup> |
| --- | --- | --- | --- | --- |
|  | Roasted<sup>c</sup> | Raw<sup>c</sup> |  |  |
| Spinach | 31 ± 1.1 | 1.0 ± 0.1 | 30 | 1 |
| Broccoli | 70 ± 0.4 | 3.3 ± 0.4 | 21 | 2 |
| Beef | 150 ± 18 | 10 ± 2.1 | 15 | 3 |
| Carrot | 17 ± 1.3 | 1.3 ± 0.1 | 13 | 4 |
| Mushroom | 65 ± 4.7 | 5.3 ± 0.6 | 12 | 5 |
| Potato | 40 ± 7.8 | 3.6 ± 0.7 | 11 | 6 |
| Garlic | 6.4 ± 0.7 | 1.0 ± 0.2 | 6.5 | 7 |
| Green bean | 16 ± 0.8 | 2.9 ± 0.2 | 5.5 | 8 |
| Salmon | 62 ± 3.2 | 11 ± 0.8 | 5.5 | 9 |
| Cauliflower | 9.4 ± 0.5 | 2.9 ± 0.3 | 3.2 | 10 |
| Pork | 25 ± 1.2 | 11 ± 1.5 | 2.3 | 11 |
| Almond | 0.7 ± 0.1 | 0.3 ± 0.03 | 2.2 | 12 |
| Asparagus | 13 ± 1.2 | 7.9 ± 1.1 | 1.7 | 13 |
| Beef substitute | 7.1 ± 0.1 | 5.8 ± 0.6 | 1.2 | 14 |
| Cheese | 8.3 ± 0.8 | 7.1 ± 0.8 | 1.2 | 15 |
| Chicken | 2.2 ± 0.1 | 2.7 ± 0.1 | 0.81 | 16 |
| Onion | 77 ± 23 | 100 ± 24 | 0.77 | 17 |
| White fish | 5.6 ± 0.2 | 7.8 ± 2.1 | 0.72 | 18 |
| Shrimp | 7.7 ± 1.3 | 15 ± 0.4 | 0.52 | 19 |
| Bacon | 0.8 ± 0.7 | 2.2 ± 0.9 | 0.36 | 20 |
| Tofu | 0.7 ± 0.02 | 2.1 ± 0.04 | 0.33 | 21 |

<sup>a</sup> Expressed as lesions per 10<sup>6</sup> DNA bases (roasted and raw). <sup>b</sup> Fold-increase (roasted/raw) calculated and ranked accordingly. <sup>c</sup>
Data represent mean ± S.D. from three replicates (*n* = 3).

**Table 4.** Dietary exposure to DNA damage (dU and 8-oxo-dG) from roasted food ingredients.

| Food Ingredients | dU Exposure<br>($\mu\text{gDNA/gFood} \cdot 10^6$ DNA bases)<sup>a, b</sup> | Rank<sup>c</sup> | 8-oxo-dG Exposure<br>($\mu\text{gDNA/gFood} \cdot 10^6$ DNA bases)<sup>a, b</sup> | Rank<sup>c</sup> |
| --- | --- | --- | --- | --- |
| Salmon | 1300000 ± 58000 | 1 | 83000 ± 4400 | 2 |
| Mushroom | 750000 ± 48000 | 2 | 310000 ± 28000 | 1 |
| Shrimp | 240000 ± 16000 | 3 | 16000 ± 2600 | 11 |
| Chicken | 200000 ± 15000 | 4 | 5300 ± 330 | 15 |
| Cauliflower | 160000 ± 14000 | 5 | 35000 ± 1800 | 5 |
| Beef substitute | 130000 ± 8500 | 6 | 16000 ± 370 | 10 |
| Beef | 120000 ± 25000 | 7 | 28000 ± 390 | 7 |
| Garlic | 110000 ± 6800 | 8 | 18000 ± 2100 | 9 |
| Cheese | 44000 ± 12000 | 9 | 1000 ± 270 | 18 |
| Onion | 41000 ± 2100 | 10 | 32000 ± 9500 | 6 |
| Broccoli | 32000 ± 2400 | 11 | 72000 ± 4100 | 3 |
| White fish | 21000 ± 530 | 12 | 2000 ± 73 | 17 |
| Green bean | 17000 ± 150 | 13 | 21000 ± 1000 | 8 |
| Spinach | 15000 ± 730 | 14 | 39000 ± 1500 | 4 |
| Pork | 13000 ± 700 | 15 | 7100 ± 390 | 13 |
| Asparagus | 8900 ± 870 | 16 | 14000 ± 1400 | 12 |
| Tofu | 8200 ± 1300 | 17 | 810 ± 91 | 19 |
| Almond | 4500 ± 350 | 18 | 260 ± 24 | 20 |
| Carrot | 1700 ± 100 | 19 | 6000 ± 470 | 14 |
| Potato | 870 ± 32 | 20 | 3800 ± 750 | 16 |
| Bacon | 230 ± 47 | 21 | 78 ± 69 | 21 |

<sup>a</sup> Exposure calculated as DNA damage (lesions per $10^6$ DNA bases) multiplied by extractable DNA content ( $\mu\text{g DNA/g food}$ ). <sup>b</sup>
Data represent mean ± propagated S.D., based on the uncertainties of both extractable DNA and lesion levels ( $n = 3$ ). <sup>c</sup> Foods were ranked separately by dU and 8-oxo-dG exposure levels.

**Figure 3.**
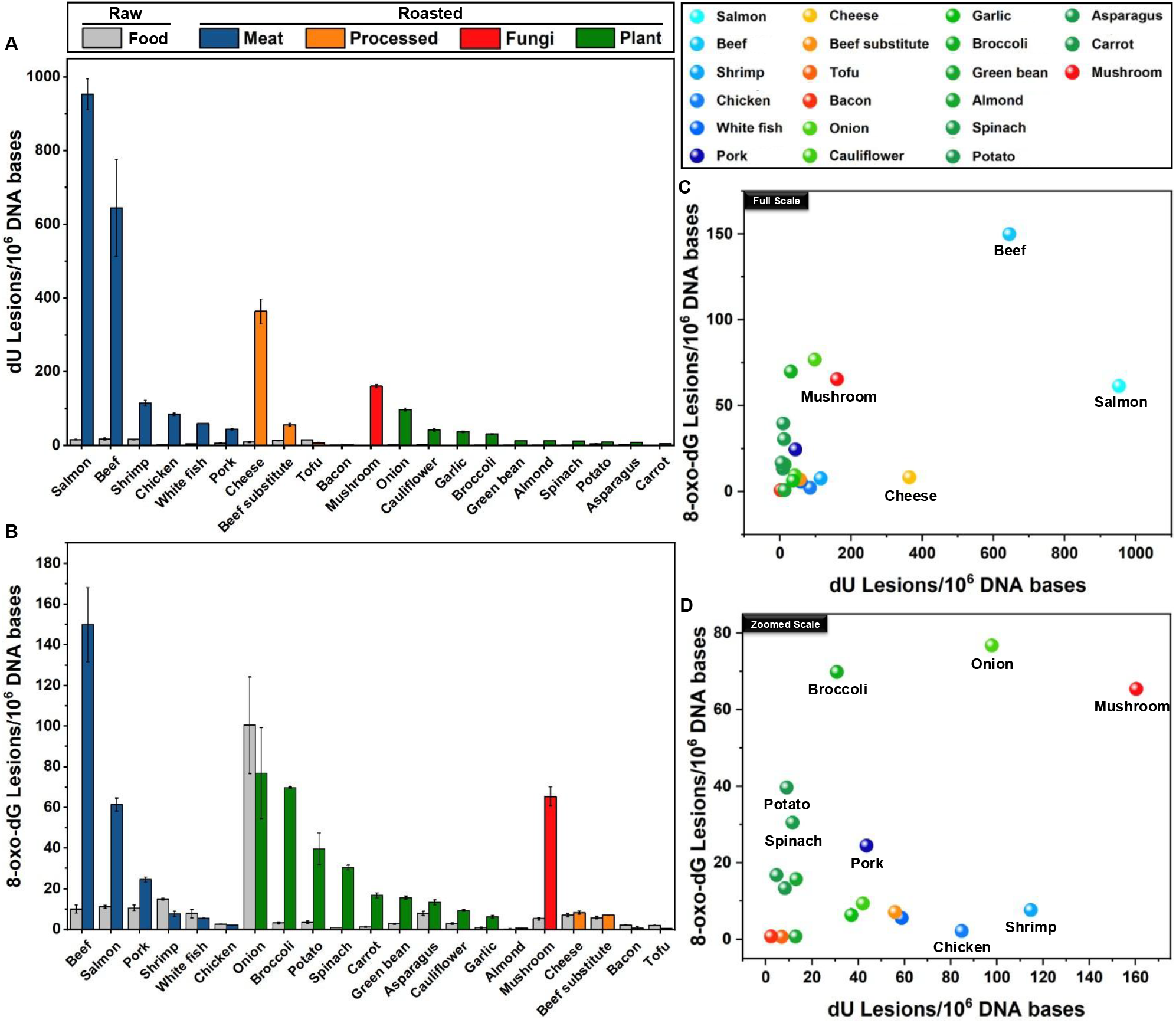
DNA damage levels in extracted DNA from raw and roasted food ingredients. (A) Levels of deaminative damage (dU), expressed as lesions per 10^6^ DNA bases. (B) Levels of oxidative damage (8-oxo-dG), expressed as lesions per 10^6^ DNA bases. Data represent the mean ± S.D. from three replicates (*n* = 3). (C) Two-dimensional correlation plot comparing dU and 8-oxo-dG damage frequencies in roasted food ingredients, illustrating differences in damage formation profiles across food categories. (D) Zoomed-in view of the correlation plot in (C), highlighting data points clustered at lower damage levels.

Fig. 3B plots 8-oxo-dG content in the raw and cooked foods (see also numerical data in Table 2). Overall, cooking increased levels of this oxidative damage in the extracted DNA in the majority (14/21) of cases, with increases of up to ≈10-fold after roasting (see fold increases in Table 3). The remainder showed no change or a moderate decrease upon roasting. In general, levels of 8-oxo-dG were several-fold lower than those for dU. Highest levels of 8-oxo-dG in roasted foods were found in beef, broccoli, mushroom, and onion, at *ca*. 50 lesions/10^6^ bases to 150 lesions/10^6^ bases. Lowest levels were measured for almond, bacon, and tofu, at 2 lesions/10^6^ bases to 4 lesions/10^6^ bases.

To gain further insights into the physicochemical origins of this damage that arises in food DNA, we asked whether levels of deaminative versus oxidative damage in the DNA extracted from heated foods are correlated. A 2D plot of damage frequencies from the 21 food sources after roasting is given in Figs. 3C and 3D. Interestingly, certain food sources (*e*.*g*., shrimp and cheese) displayed high levels of dU with very low levels of oxidative damage; conversely, other foods (*e*.*g*., potato, broccoli, and spinach) showed a propensity for high levels of oxidative DNA damage with relatively little deamination. Overall, the results reveal no discernible correlation in deaminative versus oxidative DNA damage among the different foods, reflecting different mechanisms for their formation and different biochemical settings in the varied foods. This is consistent with known mechanisms; reactive oxygen species are widely recognized to be responsible for the formation of 8-oxo-dG,^32^ whereas simple heat alone (in the presence of water) readily causes hydrolytic deamination of deoxycytidine to deoxyuridine.^31^ Nevertheless, heat likely acts as an indirect accelerant of oxidation for the former species, as documented by the general increases observed here in levels after roasting (Table 3).

The total exposure to such lesions in a diet containing these foods is expected to depend both on the frequency of the damage occurrence in a given amount of DNA and on the total extractable DNA content consumed per gram of food. Multiplying these two yields a relative measure of the food exposure in a meal to each lesion as lesions/g. In Fig. 4, these exposure values from our roasted food data are plotted. Fig. 4A shows the data for dU with the exposures ranked; results show that exposure is as high as 1.3x10^6^ lesions per gram of food, with salmon and mushrooms providing by far the greatest exposures. Lowest exposures are seen for potato and bacon, lower by 2-3 orders of magnitude than the high examples. No apparent correlation is seen between different classes of foods (*e*.*g*., meat vs. plants). For 8-oxo-dG (Fig. 4B), exposures are as high as 3.0x10^5^ lesions per gram. Greatest exposures are seen for mushrooms, salmon, and broccoli. Interestingly, seven of the ten highest exposures come from plants and fungi. Lowest exposures of this oxidatively damaged nucleotide are seen for bacon and almonds, more than three orders of magnitude reduced relative to the high-exposure foods. A few foods show low exposures to both forms of damage (*e*.*g*., carrot, almond, whitefish, and potato). Bacon is also a generally low-exposure food from these results, which may be surprising given that oxidants such as nitrites and nitrates are commonly added during processing and curing.^36^

**Figure 4.**
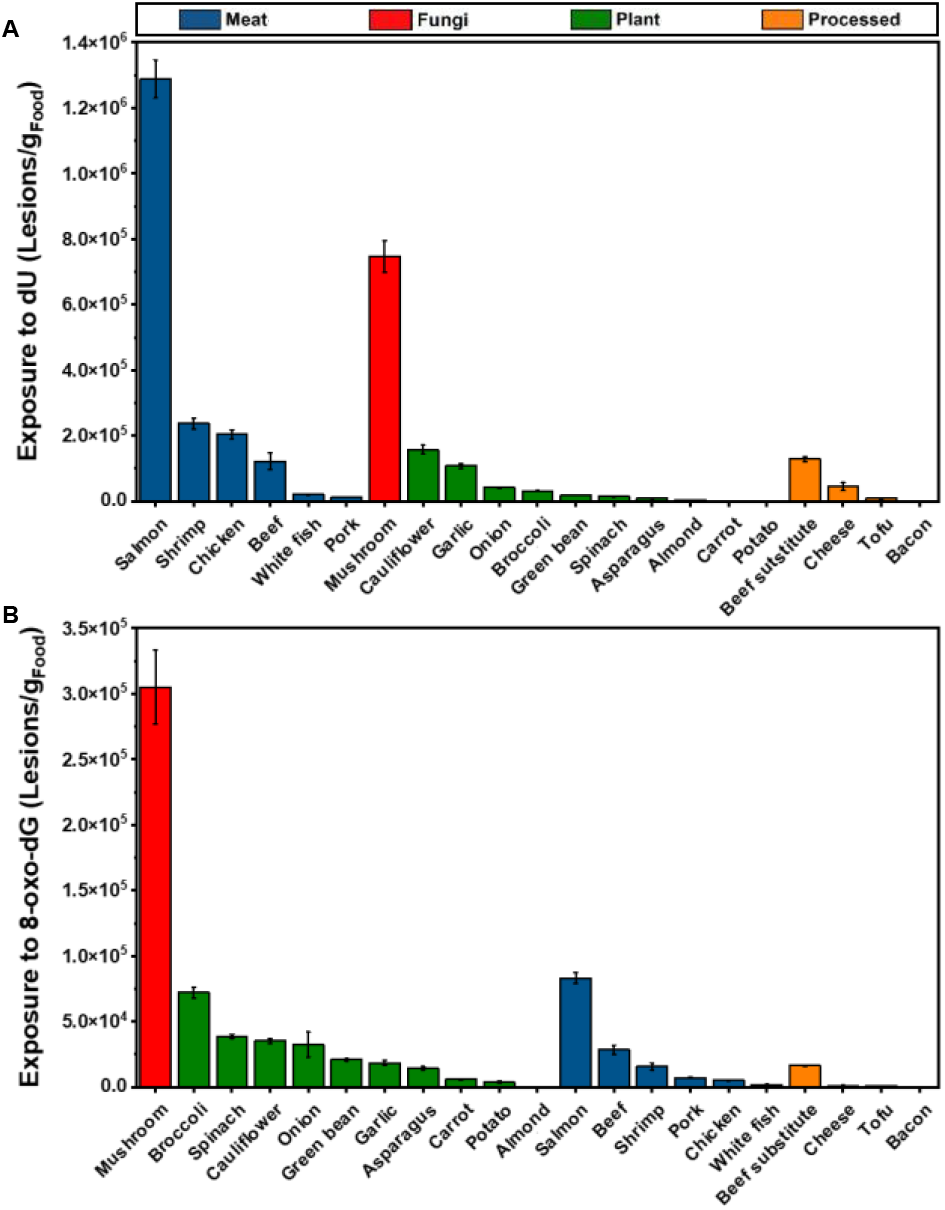
Dietary exposure levels of DNA damage resulting from consumption of roasted foods. (A) Exposure to deaminative DNA damage (dU), calculated as lesions per 10^6^ DNA bases multiplied by extractable DNA content (μg DNA per gram of food). (B) Exposure to oxidative DNA damage (8-oxo-dG), calculated similarly. Foods are categorized as meat-based, fungi-based, plant-based, or processed. Error bars represent propagated S.D. from extractable DNA and lesion level measurements (*n* = 3).

### Effects of processing temperature and method on damage content

Next, we asked how varied food preparation methods might affect DNA damage levels. Since heat levels, time, and air exposure can differ between cooking methods, this could potentially play a substantial role in the exposure to damaged DNA components.^12^ To examine this possibility, we chose five representative foods with either high damage levels or large increases upon roasting, and we compared them after varied common preparation methods: raw, roasting (15 min, 220 ℃), boiling (25 min, 100 ℃), frying (20 min, 200 ℃), and *sous vide* (60 min, 60 ℃). Conditions and methods are detailed in the Supporting file. DNA was again extracted and damage lesions (dU, 8-oxo-dG) were measured in triplicate.

Results showed widely variable DNA damage content, both among different foods and among varied cooking methods (Fig. 5). Beef, tofu, chicken, mushrooms, and salmon were tested. Overall, frying and roasting – involving the highest temperatures tested – resulted in the most elevated levels of dU (Fig. 5A), except for tofu, as previously mentioned. For 8-oxo-dG (Fig. 5B), roasting again resulted in the highest lesion levels among these preparation methods, although frying caused lesser increases than for the former lesion. For both lesions, boiling and *sous vide* methods produced the lowest damage levels, in most cases similar to those of raw foods.

**Figure 5.**
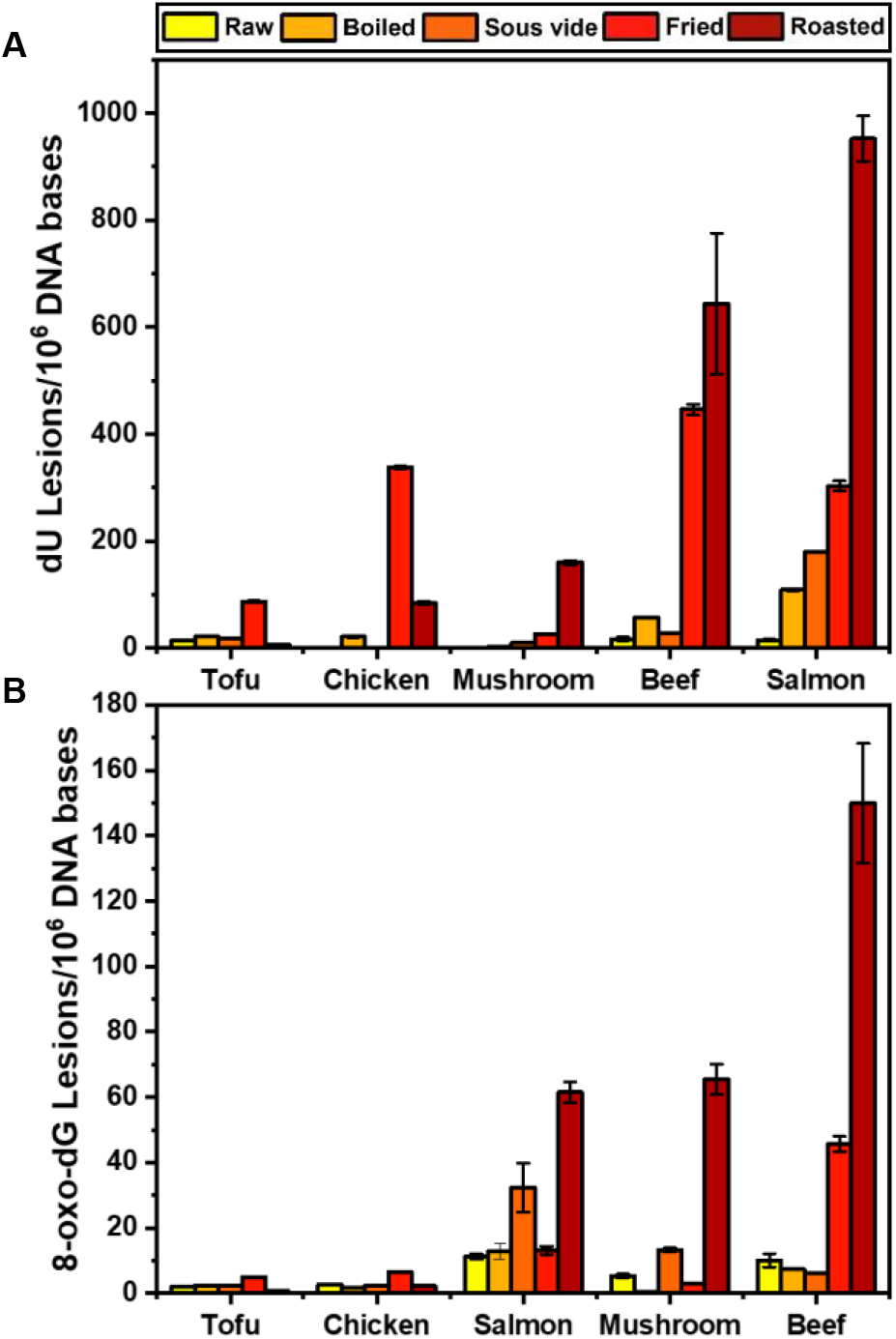
Comparison of DNA damage levels resulting from different cooking methods (raw, boiled, sous vide, fried, and roasted) in salmon, beef, and mushroom. (A) Levels of deaminative DNA damage (dU) measured as lesions per 10^6^ DNA bases. (B) Levels of oxidative DNA damage (8-oxo-dG) measured as lesions per 10^6^ DNA bases. Data represent mean ± S.D. for three replicates (*n* = 3).

### Comparing the genotoxicity of dU and 8-oxo-dG

Finally, we asked whether potential risks of metabolic uptake might be different for the two damaged nucleosides studied here, given that pyrimidine and purine salvage pathways are distinct. The presence of damaged DNA bases at high levels in genomic DNA is known to promote double-strand breaks (DSB) resulting from the accumulation of abasic sites that occur as repair intermediates from base excision repair (BER).^31^ As described above, few studies exist of the uptake of these damaged nucleosides, and none has to our knowledge tested possible genotoxicity prior to our recent report of the effects of dU. To examine this in a preliminary way, we examined the formation of DSB in HCT116 cells exposed to 8-oxo-dG or dU, detected by γ-H2AX immunostaining, a well-established marker (Figs. 6A and S1–S3). Quantification of γ-H2AX foci^37^ revealed a concentration-dependent increase in DSB for both nucleosides (Fig. 6B), with cells exposed to 1 mM dU exhibiting the highest number of γ-H2AX foci per cell (approximately 500 foci), suggesting substantial DNA damage. Interestingly, while 8-oxo-dG treatment also induced significant DSB formation, the number of γ-H2AX foci was slightly lower than in dU-treated cells at equivalent concentrations.

**Figure 6.**
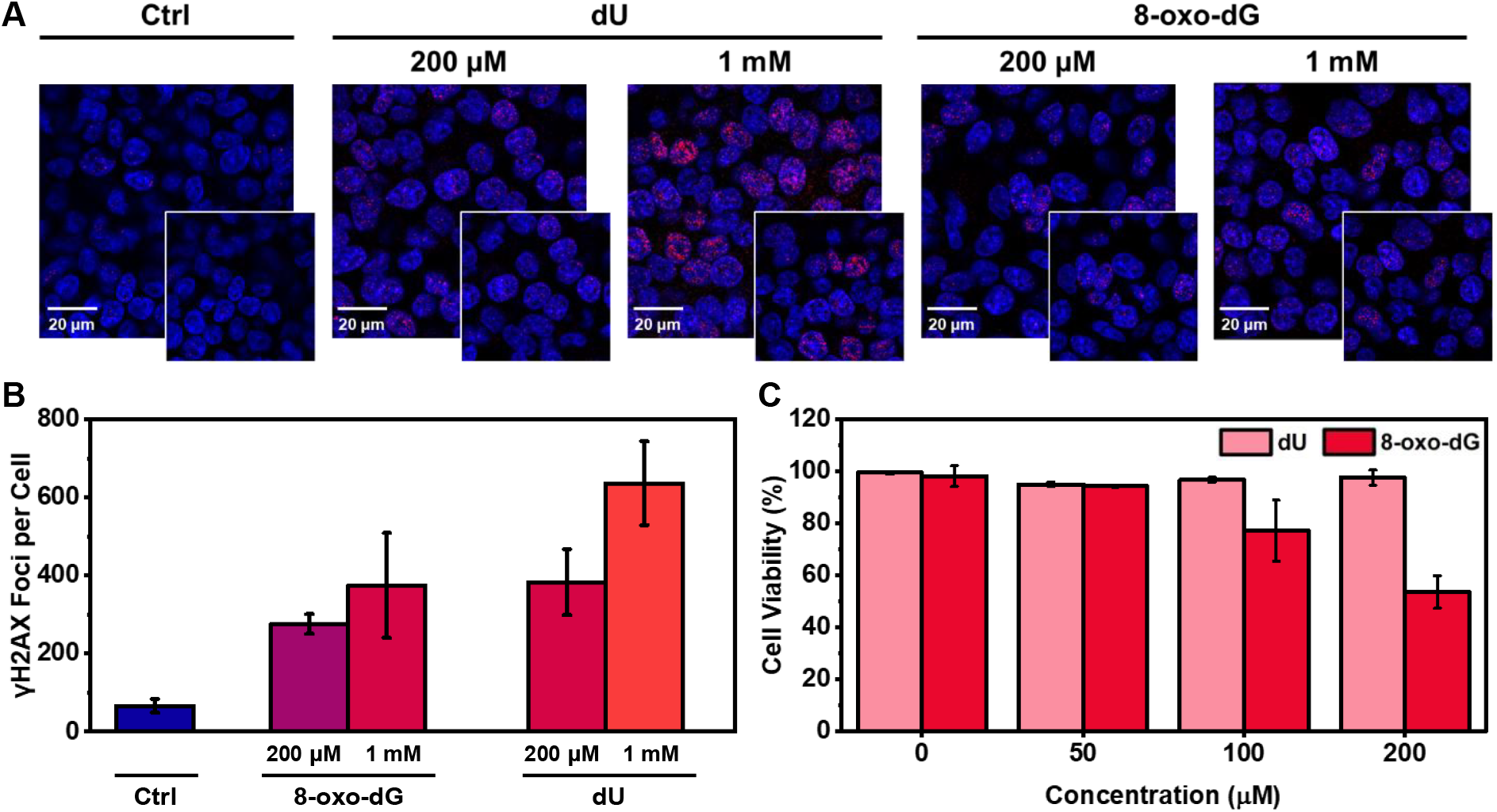
γ-H2AX immunostaining and quantification of DNA double-strand breaks (DSBs) in HCT116 cells treated with damaged nucleosides. (A) Representative confocal fluorescence images of cells stained with γ-H2AX (magenta) and Hoechst 33342 (blue) after 24 h treatment with 200 µmol/L or 1 mM of 8-oxo-dG or dU. White boxes indicate magnified insets. Scale bars = 20 μm. (B) Quantification of γ-H2AX foci per cell across treatment conditions. Data represent mean ± S.D. for four replicates (*n* = 4). (C) Cell viability treated by dU or 8-oxo-dG. Data represent mean ± S.D. for three replicates(*n* = 3).

### Differential effects on cell viability

Despite inducing fewer γ-H2AX foci, 8-oxo-dG exhibited markedly stronger cytotoxicity compared to dU (Fig. 6C). While dU treatment maintained full cell viability above 90 % even at 200 μmol/L, 8-oxo-dG caused a concentration-dependent decrease in viability, reaching approximately 50 % at 200 μmol/L. This observation contrasts with a previous report^12^ where dU demonstrated slightly higher cytotoxicity in colony formation assays.

Our study addresses how DNA in foods, which is subject to digestion and salvage into human DNA, varies both in quantity and in potentially hazardous damage levels across a variety of food types and preparation methods. DNA salvage from the diet remains poorly characterized, and it is not yet known what fraction of humans’ cellular DNA is composed of deoxynucleotides from the diet relative to those obtained by intracellular *de novo* synthesis. In addition, although partial data exist,^19^ it is not yet broadly clear which tissues make greater use of deoxynucleotide salvage relative to others, nor is it known how and whether the ratio of salvage to *de novo* pathways changes with aging. Nevertheless, it is clear that this nucleotide uptake does occur to a significant extent, and the existence of these salvage pathways in human cells suggests consideration of the amounts and quality of DNA in foods that are common in human diets. Although considerably more work is needed to better quantify this salvage, the current findings provide a broad measure of extractable DNA content in foods, and show that the levels of DNA in foods varies widely. Given that this is subject to salvage to some degree, it seems relevant to consider DNA content and quality in the diet. A study of three uncooked foods by Berdal measured levels of damaged nucleosides and considered the possibility that some of these might be incorporated into cellular DNA.^38^ A small number of prior studies have tested cellular uptake of damaged nucleotides to evaluate the effects of extracellular damage on nucleotide pools. Goulian *et al*. in 1985 measured cytotoxicity after exposure of cells to deoxyuridine, which led to an imbalance of the thymidine nucleotide pool.^39^ More recently, Mundt *et al*. incubated cell lines with radiolabeled 8-oxo-dG, and found that some of the damaged purine base was incorporated into DNA via depurination and ribonucleotide synthesis typical for purine salvage.^40^ While neither of those studies considered the possibility of salvage of nucleotides from cooked foods, they do add support to the notion that the damaged noncanonical nucleotides measured here may find their way into cellular DNA.

There appears to be relatively little information in prior literature on DNA content of foods; a literature survey found relatively few measurements of DNA content alone in earlier studies (a 1995 review cites DNA/RNA mixtures,^16^ while other studies measured DNA from foods based on dry weight).^17, 18^ In the current measurements of DNA content of fresh foods, we observe highly variable levels, indicating widely varied consumption of DNA in humans depending on their dietary choices. Based on the current findings, one 100 g serving of roasted mushrooms implies consumption of ca. 460 mg of extractable DNA, whereas 100 g of roasted carrots yields only 36 mg. Some of this variation could well come from distinct matrices from diverse foods that render DNA more or less accessible to our extraction method. This variable accessibility almost certainly also plays a role in how humans are exposed to DNA in foods, as the matrices of different foods likely release DNA differentially during chewing and digestion. Much of the variability that we observe in DNA content may also reflect biological differences in the food type; for example, onion (which has low extractable DNA content) has very large cells and thus fewer nuclei per gram of food, while fungi have considerably smaller cells and thus mushrooms likely have considerably more nuclei per gram.^41^ The sizes of genomes also could potentially play a role; for example, plant genomes are often considerably larger than mammalian genomes, and thus potentially provide greater DNA content;^42^ however, we do not observe a general trend toward greater extractable DNA content in plants tested here. Other factors that may play roles are variable levels of DNA from mitochondria and chloroplasts. Overall, our measurements suggest that common human diets likely involve consumption of 1 g to 2 g of DNA per day, consistent with prior estimates,^16^ and perhaps more if extraction via chewing and digestion is more efficient than our protocol.

Our results show, in addition, that cooking of foods markedly increases levels of damaged nucleotides that are ingested. For deoxyuridine, we find that mild roasting can increase levels by a factor of up to 250-fold. We find dU content as high as 950 lesions per million bases after cooking; given an assumption of 1.5 g/day of total DNA in a typical diet, this implies exposure of up to 1.5 milligrams per day of this damaged nucleoside. We have hypothesized that certain damaged deoxynucleotides, such as deoxyuridine, may be metabolically salvaged to a significant extent by cells and tissues, as first documented for dU in earlier studies.^12^ Preliminary data suggest that pyrimidines such as dU can be incorporated into intracellular DNA in cultured cells upon simple incubation, yielding measurable damage and DNA damage and repair responses. In addition, a test of feeding deoxyuridine to mice resulted in measurable increases in DNA repair responses and in up to 15-fold increased quantities of dU content in intestinal genomic DNA.^12^ We suggest that, given the knowledge that salvage of modified nucleosides is widely employed for metabolic labeling experiments and in anticancer therapies, it should not be particularly surprising if certain damaged nucleotides, especially pyrimidines, might also be metabolically incorporated. However, much more work is needed to examine these and other forms of damaged nucleosides before broad claims or risks assessments can be made.

Given that caveat, the current work helps to highlight the variable levels of damaged DNA exposure that human foods present, and documents how food choices and cooking methods may increase human exposure to these components. This may be particularly relevant for clinical populations that are genetically predisposed to cancer of the digestive tract. For example, populations with Familial Adenomatous Polyposis (FAP) acquire colorectal cancers at very high rates due to a deficiency in the APC tumor suppressor,^43^ and avoiding exposure to agents in foods that may promote DNA damage seems warranted. Recent studies aimed at benefiting FAP patients have suggested the adoption of low-inflammatory diets and avoidance of processed foods;^44^ the current results also suggest that DNA content and DNA damage exposure might also be factors that are worthy of future consideration.

To the extent that damaged DNA nucleotides may be subject to salvage, it is of interest to examine total levels of damaged lesions per gram of food (here denoted as exposure in lesions per gram). For damage exposure, assessed by DNA content multiplied by damage frequency, heat, time, and air exposure appear to be the chief variables that increase this exposure. Our data here show that foods vary widely in lesion exposure, both because they vary widely in DNA content and because the DNA in disparate foods is affected very differently by cooking. This wide variability in damage exposure – over three orders of magnitude - suggests that the biochemical context for each food, as well as cooking methods, can make a large difference in the damage that occurs to the DNA. It will be of interest in the future to explore physical, enzymatic, or chemical components of certain foods that appear to protect the contained DNA from damage on heating.

We observed that the deaminated nucleoside dU displays contrasting behavior between DSB formation and cell viability in HCT116 cells (Fig. 6), showing elevated damage levels but limited toxicity. Although dU exposure generates numerous DSBs, these breaks are predominantly repaired through homologous recombination or non-homologous end-joining pathways.^45^ The efficiency of these repair mechanisms may explain why dU-treated cells maintain relatively high viability despite substantial DNA damage. However, more extended exposure (see below) may lead to toxicity, and DSB can also lead to mutagenesis when errors in end-joining repair occur.

While we observed no toxicity of dU after 24 h with the HCT116 cell line, a previous study reported significant cytotoxicity in Chinese Hamster Ovary (CHO) cells for that nucleoside.^12^ The temporal differences between our short-term viability assays and previously reported colony formation assay (carried out over several days) may contribute to the observed variance in outcomes. While γ-H2AX foci represent an immediate cellular response to DNA damage, colony formation assesses long-term survival and proliferation capacity. Different cell types, experimental conditions, and assay endpoints could further influence these results.

In living form, plant and animal food sources have active DNA repair enzymes that continuously remove these damaged components,^46, 47^ and freshly harvested raw foods may continue this repair to some extent as enzymes presumably remain active temporarily. Consistent with this notion, we find that raw foods have relatively undamaged DNA, and exposure to damaged nucleotides in human diets could in principle be minimized via consumption of raw foods when possible. However, a number of foods are rarely eaten raw (*e*.*g*., poultry, pork, and beef substitute), and our results show that choice of cooking methods for such foods can make a large difference in this exposure. While the current experiments begin to quantify this exposure in a range of dietary components, more work is needed to evaluate any risks resulting from it.

## AUTHOR INFORMATION

### Author

Jinwoo Shin − Department of Chemistry, Stanford University, Stanford, CA 94305, USA.

Pawel Jaruga − Biomolecular Measurement Division, National Institute of Standards and Technology, Gaithersburg, Maryland 20899, USA.

±(deceased on Dec 13, 2024) Miral Dizdaroglu −Biomolecular Measurement Division, National Institute of Standards and Technology, Gaithersburg, Maryland 20899, USA.

### Author Contributions

J.S. performed DNA extraction from food samples and conducted the biological assays. P.J. and M.D. quantified DNA damage in the extracted samples. E.T.K. conceived and designed the project. J.S. and E.T.K. wrote the manuscript. E.T.K. provided project supervision and manuscript editing.

### Notes

The authors declare no competing financial interest. The graphical abstract and figures were created in BioRender. Shin, J. (2025) https://BioRender.com/ozwms6b

### NIST Disclaimer

Certain equipment, instruments, software, or materials are identified in this paper in order to specify the experimental procedure adequately. Such identification is not intended to imply recommendation or endorsement of any product or service by NIST, nor is it intended to imply that the materials or equipment identified are necessarily the best available for the purpose.

## Table of Contents (TOC)

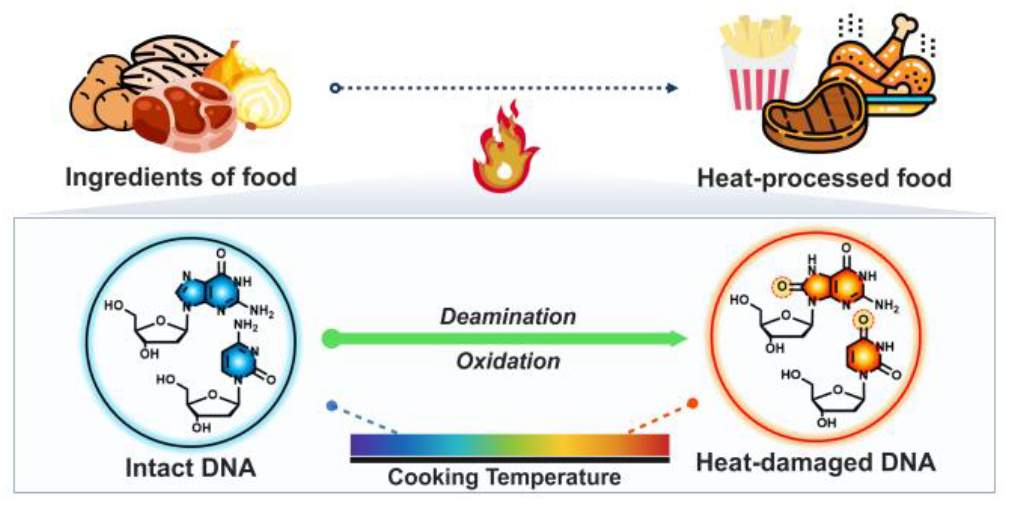

