## Supporting Information for "DNA Content and DNA Damage in Raw and Heat-processed Foods"

#### Materials and Methods

##### Instrumentation

DNA quantification extracted from food ingredients is measured using a Thermo Scientific Invitrogen Nanodrop One Spectrophotometer (13-400-525). Fluorescence images were recorded on an inverted Zeiss LSM 780 multiphoton laser scanning confocal microscope in the Cell Science Imaging Facility (CSIF) at Stanford. Products of DNA damage were identified and quantified by GC-MS/MS and LC-MS/MS with isotope dilution using stable isotope-labeled analogues of the products as internal standards. Luminescence was measured using a Thermo Scientific Fluoroskan Ascent.

##### Materials

Food ingredients were purchased from Safeway (Menlo Park, CA, USA). Easy-DNA gDNA purification kit was purchased from Thermo Fisher Scientific (K180001). Damaged nucleosides used for cellular experiments were purchased from Sigma-Aldrich. Phospho-Histone H2AX (Ser 139) Rabbit mAb was purchased from Cell Signaling Technology (9718S). This was visualized with Goat anti-Rabbit IgG secondary antibody, Alexa Fluor Plus 647 (Invitrogen, A32733). 2′-Deoxyuridine-$^{13}C_9$,$^{15}N_2$ and 8-oxo-dG-$^{15}N_5$ were purchased from Cambridge Isotope Laboratories, Inc. (Tewksbury, MA, USA

##### DNA extraction from foods

Food ingredients were cut into cubes (1 cm × 1 cm) prior to cooking and prepared using different cooking methods (raw, boiling, *sous vide*, frying, or roasting). For the boiling process, each food was boiled in boiling water (ca 100 ℃) for 25 min. For the sous vide process, each food was vacuum-sealed and cooked in a sous vide device set at 60 ℃ for 1 h. For the frying process, each food was cooked in a preheated pan containing oil maintained at 200 ℃ (monitored by thermometer) for 20 min, turning the food every 5 min to ensure even cooking. For the roasting process, a standard kitchen oven was pre-heated to 220 °C, and food samples were roasted for 15 min. An Easy-DNA gDNA purification kit was used for extracting DNA from food samples. Briefly, food samples were dried thoroughly with paper towels to remove excess moisture, chopped into small pieces, and incubated with solution A and solution B (provided in the kit) according to the manufacturer’s protocol. After chloroform extraction and centrifugation, DNA was precipitated and purified by ethanol purification following the manufacturer's instructions.

**Quantification of DNA damage in extracted DNA**

Aliquots of 50 µg of three independently isolated DNA samples from each beef, pork and potato samples were used for LC-MS/MS measurementsof 8-oxo-dG and dU using 8-oxo-dG-$^{15}N_5$, and dU-$^{13}C_9$,$^{15}N_2$ as their internal standards, respectively. Hydrolysis of DNA samples to release these products from DNA as nucleosides, and other measurement conditions have been described in detail elsewhere. The mass/charge transitions used for these measurements were *m/z* 229 → *m/z* 113 for dU, *m/z* 240→ *m/z* 119 for dU-$^{13}C_9$,$^{15}N_2$, m/z 284→m/z 168 for 8-oxo-dG, and m/z 289→m/z 173 for 8-oxo-dG-$^{15}N_5$.

Cytotoxicity was calculated using the following formula:

$$\%\ \text{Cytotoxicity} = 100 \times \frac{\text{Sample LDH Release} - \text{Negative Control}}{\text{Maximum LDH Release} - \text{Negative Control}}$$

Cell viability was determined as:

$$\%\ \text{Cell Viability} = 100 - \%\ \text{Cytotoxicity}$$

### Supporting Figures and Tables

**Table S1.** Food ingredients and sources

| Food Ingredients | Type or cut | Source |
|---|---|---|
| **Meat-based** | | |
| Beef | Ground (20 % fat and 80 % lean) | ASPEN RIDGE |
| Chicken | Boneless skinless breast tenders | Organic |
| Pork | Ground (20 % fat and 80 % lean) | OPEN NATURE |
| Salmon | Ground (20 % fat and 80 % lean) | Safeway |
| Shrimp | Breast | Safeway |
| White Fish | Dover | Safeway |
| **Processed-derived** | | |
| Bacon | Raw, cured | HEMPLER'S |
| Beef Substitute | Beyond beef | Beyond Beef |
| Cheese | Cheddar | Safeway |
| Tofu | Firm | Safeway |
| **Plant-based** | | |
| Almond | Raw, whole | Safeway |
| Asparagus | - | Safeway |
| Broccoli | - | Safeway |
| Carrot | - | Safeway |
| Cauliflower | - | Safeway |
| Garlic | Fresh peeled | Christopher Ranch |
| Green bean | - | Safeway |
| Onion | Yellow | Safeway |
| Potato | White | Safeway |
| Spinach | - | Safeway |
| **Fungi-based** | | |
| Mushroom | - | Safeway |

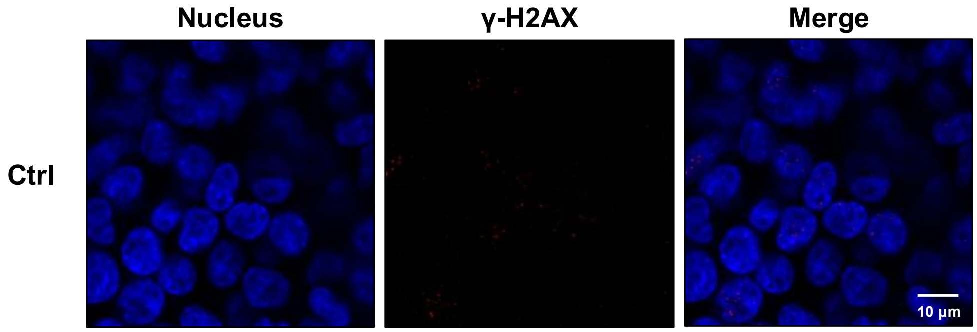

**Figure S1. γ-H2AX immunostaining of DSB in HCT116 cells without damaged nucleoside treatment.** Representative confocal fluorescence images of cells stained with γ-H2AX (magenta) and Hoechst 33342 (blue). Scale bars = 10 µm.

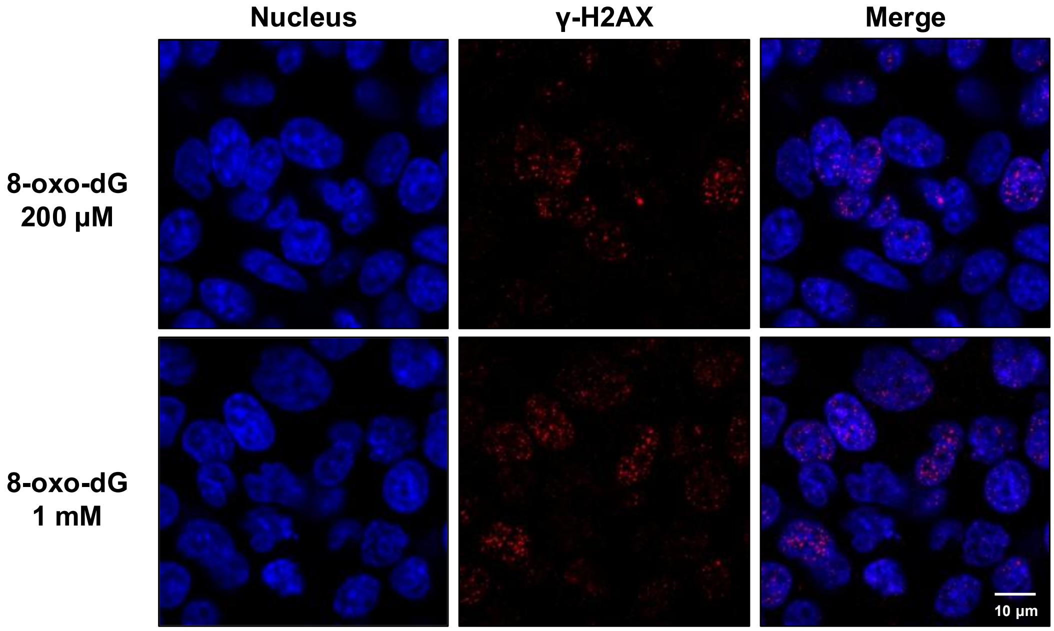

**Figure S2. γ-H2AX immunostaining of DSB in HCT116 cells incubated with the oxidatively damaged nucleoside 8-oxo-dG.** Representative confocal fluorescence images of cells stained with γ-H2AX (magenta) and Hoechst 33342 (blue). Scale bars = 10 µm.

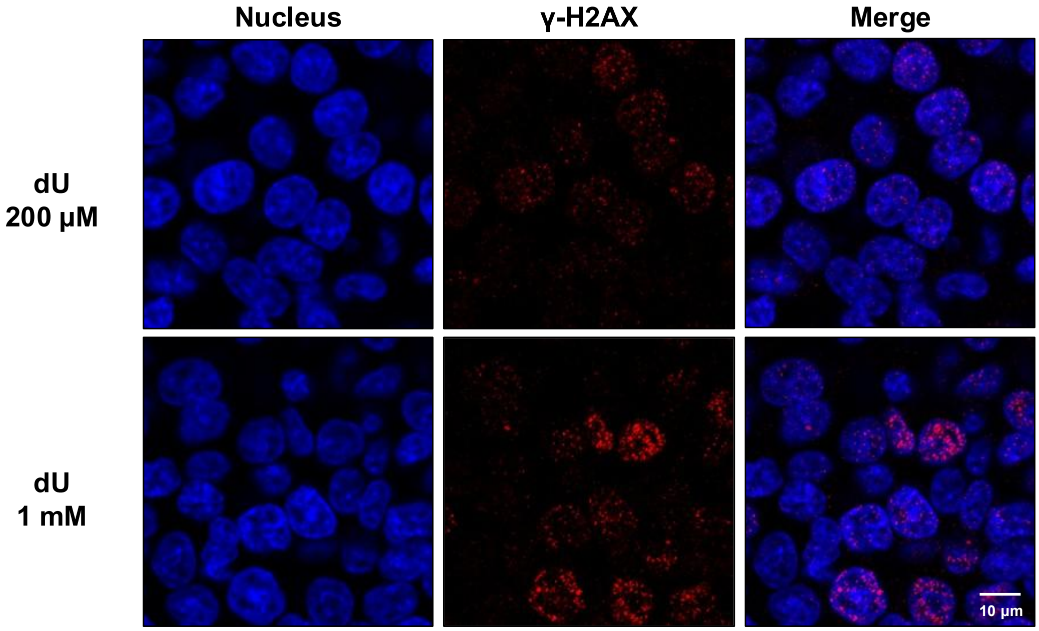

**Figure S3. γ-H2AX immunostaining of DSB in HCT116 cells treated with the deaminated nucleoside dU.** Representative confocal fluorescence images of cells stained with γ-H2AX (magenta) and Hoechst 33342 (blue). Scale bars = 10 μm.

**NIST Disclaimer**

Certain equipment, instruments, software, or materials are identified in this paper in order to specify the experimental procedure adequately. Such identification is not intended to imply recommendation or endorsement of any product or service by NIST, nor is it intended to imply that the materials or equipment identified are necessarily the best available for the purpose.
